# *In Silico* design and characterization of multi-epitopes vaccine for SARS-CoV2 from its spike proteins

**DOI:** 10.1101/2020.06.03.131755

**Authors:** Gunderao H Kathwate

## Abstract

COVID 19 is disease caused by novel corona virus, SARS-CoV2 originated in China most probably of Bat origin. Till date, no specific vaccine or drug has been discovered to tackle the infections caused by SARS-CoV2. In response to this pandemic, we utilized bioinformatics knowledge to develop efficient vaccine candidate against SARS-CoV2. Designed vaccine was rich in effective BCR and TCR epitopes screened from the sequence of S-protein of SARS-CoV2. Predicted BCR and TCR epitopes were antigenic in nature non-toxic and probably non-allergen. Modelled and refined tertiary structure was predicted as valid for further use. Protein-Protein interaction prediction of TLR2/4 and designed vaccine indicates promising binding. Designed multiepitope vaccine has induced cell mediated and humoral immunity along with increased interferon gamma response. Macrophages and dendritic cells were also found increased over the vaccine exposure. *In silico* codon optimization and cloning in expression vector indicates that vaccine can be efficiently expressed in *E. coli*. In conclusion, predicted vaccine is a good antigen, probable no allergen and has potential to induce cellular and humoral immunity.

## Introduction

In December 2019 group of patients from Wuhan city of China was found to have pneumonia like symptoms and diagnosed with the infection of beta coronavirus(1). Later it spread across the china and now spread all over the globe. These infections were named as COVID 19 disease and the virus as SARS-CoV2 by WHO(2). On 30 January 2020 due to its spread more than 120 countries declared COVID 19 pandemic as a public health emergency of international concern. Novelty of SARS-CoV2 is its rapid spread may be due asymptomatic patients(3,4) and highly sophisticated, time consuming diagnostic methods(5–7). As of 13 May 2020, 7 million confirmed cases with more than 400,000 deaths worldwide(8). In India due to lockdown positive cases are in control and spread is also marginal, but still after approximately four months’ case reports are increases. As of 13 May 2020, a total of 71865 confirmed cases with 2415 deaths are reported by ministry of health and family welfare, Govt. of India. Till date, no antiviral drugs available to combat the SARS-CoV2 infections. There are few drug candidates in clinical trials but the process is time consuming(9). In South Korea, Lopinavir/Ritonavir combination was found significantly effective in lowering of viral load to no detectible or little SAR-CoV2 titer(10). But in another group study same combination was totally ineffective beyond standard care(11). Another drug pair drug, hydroxychloroquine, an antimalarial drug and azithromycin reported to found effectively associated with reduction of viral load(12,13). But QT interval prolongation that may cause life threating arrhythmia(14) and in large scale study both drugs and drug alone compared with neither drugs was not associated with mortality(15,16). In a drug repurposing study, remdesivir, lopinavir, emetine, and homoharringtonine have significantly inhibited replication of SAR-CoV2(17). But this was *in vitro* study, clinical trial reports are underway. Similar determined attempts are going through the globe. To break the chain of infection prevention is much better option than treatments. Several evidences suggest and support the importance of vaccine to eliminate COVID19 epidemic (18–20). Various epidemiological surveys provide the evidences that naturally acquired immunity can eliminate the SARS-CoV2 titer(21,22). More than 80% patients develop mild and 14% develop severe symptoms of COVID19 with zero case fatality rate(23).

Presently there is no vaccine available against COVID19 disease. Efforts are being taken for the development of effective vaccine all over the globe(24–27). All these vaccines are under development(28) and may require at least 1to 1.5 year to hit the market(29). Bioinformatics approach can accelerate the process by predicting potential candidate peptides. Vaccines against MERS, Ebola, and human papilloma virus were designed and successfully developed by using bioinformatics approaches(30).

SARS-CoV2 belong to beta coronoviridae family which includes four endemic viz HCoV-HKU1, HCoV-OC43, HCoV-229E, HCoV-NL63, two epidemic viruses like Middle East Respiratory Syndrome virus (MERS), and SARS(31). This is a non-segmented positive strand RNA virus with envelop. Envelop proteins are categorized into structural, non-structural and accessory proteins. Structural proteins are involved in protection and bind to the host. Several proteins of SAR-CoV2 are important for virulence and pathogenesis. For example Nucleocapsid (N) protein N, is essential for RNA binding and its replication and transcription(32). Envelope (E) and membrane (M) proteins and instrumental in virus assembly and virulence promotion(33,34). E and M proteins are effective in induction of immune response(24). Spike (S), a structural protein is responsible for binding to receptor on host cell, Angiotensin converting enzyme 2(35). Thereby proteolytic cleavage by TMPRSS2 allows subsequent entry into cell through endocytosis(36). S protein also found to be involved in activation of T cell response(37,38). Entire S protein expressed in chimpanzee adeno 38 (ChAd)-vector showed protection against SARS-CoV2 in mice and *rhesus* macaques(39). Single dose of ChAdOx1 nCoV-19 vaccine is sufficient to elicit humoral and cell mediated immune response in both animals. Viral load was found reduced compared to control animals and symptoms of pneumonia were absent. Here we designed a multi-epitopes vaccine from S Protein. This vaccine has all the ideal properties like stability at room temperature, immunogenic, antigenic and non-allergen. All the epitopes are good in stimulation of humeral as well and cell mediated immunity.

## Methods

### Selection of protein for epitope prediction

Complete Spike protein sequence (P0DTC2) of SARS-CoV2 was downloaded in fasta format from UniProt protein database. This sequence was used for further analysis and obtaining potential epitopes for B and T cell receptors.

### T cell Epitope prediction

There are various tools used to predict the epitopes that could be presented to T cell receptors. Potential epitopes of various bacteria and viruses are predicted by using such online epitopes predicting tools. Following tools were used for prediction of TCR (T cell Receptor) epitopes IEDB MHC-I processing predictions, MHC-NP, netCTLpan1.1, RANKPEP, and netMHCpan4.0. All these tools were used to screen the potential peptides as epitopes for TCR. These tools have various methods to predict the epitopes. IEDB (Instructor/Evaluator Database) is a web-based epitope analysis resource includes tools for T cell epitope prediction, B cell epitope prediction and other analysis tools like epitope conservancy etc. This resource use methods based on artificial neural network (ANN), stabilized matrix method (SMM), and Combinatorial Peptide Libraries(CombLib), predicts the peptides way that processed naturally and presented by MHC I (40).

### Analysis of immunological properties

Immunogenicity, Toxicity, Allergen, Conservancy, and Antigenicity were analyzed for predicted TCR epitopes. IEDB MHC class I immunogenicity server and conservancy tool were used for determination of immunogenicity and conservancy. Immunogenicity of a peptide MHC complex (http://tools.immuneepitope.org/immunogenicity/) were assessed keeping all the parameters at default. Protein sequence variants(40) used setting sequence identity 100% and other parameter default. ToxiPred (http://www.imtech.res.in/raghava/toxinpred/index.html) online tool predicts the toxicity considering physicochemical properties of selected peptides(41). Online server AllergenFP v.1.0 (http://ddgpharmfac.net/AllergenFP/) was used to predict peptides as allergens(42). Antigenicity of epitopes was analyzed by online server VaxiJen v2.0 (http://www.ddg-harmfac.net/vaxijen/VaxiJen/VaxiJen.html). Threshold value was set to 0.5 (43). This is alignment independent predictor based on auto-cross covariance (ACC) transformation epitopes sequences into uniform vectors of principle amino acid properties. Accuracy of this server varies in between 70 to 89% depending on targeted organism.

### Linear B cell receptor epitope prediction

Six different method were used for the prediction of B cell receptor (BCR) epitopes. All these methods generate fragments the protein. For the prediction of B cell epitopes, it is necessary to find linear sequence of B cell epitope in the protein sequence. BepiPred linear epitope predication server (http://www.cbs.dtu.dk/services/BepiPred/) uses a hidden Markov model and propensity scale method. Similarly, other properties are also being consider to predict good B cell epitopes(44). Those properties are calculated by different methods at IEDB server (http://tools.iedb.org/bcell/)like Kolaskar–Tongaonkar antigenicity scale provide physiology of the amino acid residues(45), Emini Surface accessible score for accessible surface of the epitope(46), Secondary structure of epitopes also has role in antigenicity. Karplus-Schulz flexibility score(47) and Chou-Fasman β turns methods(48) were used for flexibility and β turns prediction respectively. Parker hydrophilicity prediction method (49)was used for determination of *in silico* hydrophilicity of peptides.

### Engineering of multi epitope vaccine sequence

High scored and common peptides predicted by various tools were selected for deriving sequence of potential vaccine candidate. Different epitopes for T cell and B cell receptors were linked together by GPGPG and AAY. To enhance the immunogenicity, OmpA (GenBank: AFS89615.1) protein was chosen as adjuvant and was linked through EAAAK at N terminal site of the vaccine.

### Prediction of immunogenic properties of designed vaccine

Antigenicity of chimeric vaccine was predicted by VaxiJen v2.0 (http://www.ddg-pharmfac.net/vaxijen/VaxiJen/VaxiJen.html) and ANTIGENpro. VaxiJen 2.0 is alignment free antigenicity prediction tool utilizes auto cross covariance transformation of protein sequence into uniform vector vectors of principal amino acid properties(43,50). ANTIGENpro (http://scratch.proteomics.ics.uci.edu/) is also online tool utilizes protein microarray dataset for the prediction of antigenicity. Based on cross validation experiment accuracy of this server is estimated to be around 76 % using combined dataset. AllerTop v2.0 and AllergenFP were two online tools utilized for the prediction of allergenicity of chimeric protein. Amino acid E-descriptors, auto- and cross-covariance transformation, and the k nearest neighbours (kNN) machine learning methods are basis of AllerTop v2.0 (http://www.ddg-pharmfac.net/AllerTOP) for allergenicity prediction(51). AllergenFP, is descriptor-based fingerprint, alignment free tool for allergenicity prediction. The tool use four step algorithm. In first step, proteins are described based on their properties like hydrophobicity, size, secondary structures formation and relative abundance. In subsequent step, generated strings are converted into vectors of equal length by ACC. Then vectors are converted into binary fingerprints and compared in terms of the Tanimoto coefficient. Applying this approach to known allergen and non-allergens can identify the 88 % of allergen/non-allergen with Mathew’s Correlation coefficient of 0.759 (42).

### Prediction of solubility and Physiochemical properties

For Solubility prediction of multi-epitopes chimeric vaccine, PROSO II server (http://mbiljj45.bio.med.uni-muenchen.de:8888/prosoII/prosoII.seam) was utilized(52). PROSO II server works on an approach of classifier which utilizes difference between TargetDB and PDB and insoluble proteins of TargetDB. The accuracy of this server is 71% at default threshold 0.6. Protoparam, online web server was exploited for evaluation of physiochemical properties. The properties like amino acid composition, theoretical pI, instability index, in vitro and in vivo half-life, aliphatic index, molecular weight, and grand average of hydropathicity (GRAVY) were evaluated.

### Secondary and tertiary structure prediction

PSIPRED and RaptorX Property online servers were used to determine secondary structure of predicted vaccine. PSIPRED is publicly available webserver, includes two feed forward neural networks works on output obtained from PSI-BLAST. PSIPRED 3.2 attains average Q3 score of 81.6% obtained using very stringent cross approval strategies to assess its performance. RaptorX property is another web based server prediction secondary structure of protein without template(53). This server utilizes DeepCNF (Deep Convolutional Neural Fields), a new machine learning model that predicts secondary structure and solvent accessibility and disorder regions simultaneously(54). It accomplishes Q3 score of approximately 84 for 3 state secondary structure and approx. 66% for 3 state solvent accessibility.

Tertiary structure of multi-epitopes chimeric vaccine candidate was built using I TASSER online server (https://zhanglab.ccmb.med.umich.edu/I-TASSER/). The I-TASSER (Iterative Threading Assembly Refinement) web server utilize sequence-to-structure-to-function paradigm to build protein structure(55). It is top ranked 3D protein structure web server in community wide CASP experiments(56).

### Tertiary structure refinement and validation

Two web bases servers were used to refine the 3D structure of multi-epitopes chimeric protein. Initially Modrefiner server (https://zhanglab.ccmb.med.umich.edu/ModRefiner/) and finally GalaxyRefine server (http://galaxy.seoklab.org/cgi-bin/submit.cgi?type=REFINE) was used. Modrefiner server is an algorithm for atomic level structure refinement utilizes c alpha trace, main chain or atomic model. Output structure is refined in term of accurate position of side chains hydrogen bon network and less atomic overlaps(57). On other hand Galaxy server rebuild the 3D structure, performs repacking, and uses molecular dynamic simulations to accomplish overall protein structure relaxation. Structure refined by Galaxy server of best quality in accordance with community-wide CASP10 experiments(58).

ProSA-web (https://prosa.services.came.sbg.ac.at/prosa.php), The ERRAT server (http://services.mbi.ucla.edu/ERRAT/) and RAMPAGE (http://mordred.bioc.cam.ac.uk/~rapper/rampage.php) web servers were utilized for 3D structure validation obtained after galaxy server refinement(59). ProSA web server calculate the quality score as Z score that should fall in a characteristic range(60). Z score obtained for a specific input protein is in context with protein structure available in public domain. ERRATA server analyses non bonded atom-atom interaction the refined 3D structure of protein compared to high resolved crystallographic protein structures(61). Ramachandran plot displays energetically allowed and disallowed dihedral psi and phi angles of amino acids. The plot is calculated based on van der Waal radius of the protein side chain. RAMPAGE server determines Ramachandran plot for a protein that include percent residues in allowed and disallowed regions(62).

### Discontinuous B cell epitope prediction

More than 90% B cell epitopes are discontinuous that is they are present in small segments on linear protein and the brought to proximity while protein folding. Discontinues B cell epitopes for designed vaccine were predicted by Ellipro online tool (http://tools.immuneepitope.org/tools/ElliPro) at IEDB. In this tool three algorithms implemented to determine the discontinuous B cell epitopes. 3D structure of input protein is approximated as number of ellipsoid shapes, calculate protrusion index (PI) and clusters neighboring residues. Ellipro defines PI score of each residue based on the center of mass of residue residing outside the largest possible ellipsoid. In consideration with other epitope predicting tools Ellipro gave an AUC value of 0.732, best of all(63,64).

### Protein-protein docking of designed vaccine with TLR2 and TLR4

Molecular docking server HADDOCK (https://bianca.science.uu.nl/haddock2.4/) was used to see the interaction of designed vaccine and TLR4. HADDOCK (High Ambiguity Driven protein-protein DOCKing) is information derived flexible docking server(65). Galaxyrefine 3D model of multi-epitopes chimeric proteins, adjuvant TLR2 (PDB ID: 6NIG) and TLR4 (PDB ID: 4G8A) were uploaded for docking at HDDOCK server keeping all the parameter default. Finally, top five models were downloaded and by PRODIGY (PROtein binDIng enerGY prediction) webserver (https://nestor.science.uu.nl/prodigy/) was utilized for prediction of binding affinities(66).

### Immune simulation

*In silico* cloning of vaccine construct was predicted by C-ImmSim server (http://150.146.2.1/C-IMMSIM/index.php). C-immSim Server is freely available web based server utilizes position specific scoring matrix (PSSM) for the prediction of immune epitopes and machine learning techniques for immune interaction(67).

JCat (Java Codon Adaptation Tool) server was utilized for reverse translation and codon optimization. Codon optimization is carried out in order to express the construct in *E. coli* host. JCat (http://www.prodoric.de/JCat) output display sequence of nucleotide, other important properties of sequence that includes codon adaptation Index (CAI), percent GC content essential for to assess the protein expression in host(68). Finally, vaccine construct was cloned in pET30a (+) plasmid vector by adding XhoI and XbaI restriction sites at C and N terminus, respectively. Snapgene tool was used to clone the construct to ensure the cloning and expression(69).

## Results

### Prediction of B and T cell receptor Epitopes

#### T cell receptor epitopes

IEDB recommend, MHC-NP, netCTLpan1.1, RANKPEP and netMHCpan3.0 server predicted the potential candidate epitopes from the spike glycoprotein sequence of SARS-CoV2. Epitopes commonly predicated by at least four prediction tools were selected further for analysis parameter. There were four such epitopes with high score and predicted by four different tools (Table 1A). All the four epitopes were immunogenic and antigenic in nature, conserved 100 %, and predicted to be non-allergen (Table 1B). Immunogenicity scores for the predicted epitopes ranges from 0.3858 to 0.0961. All the epitopes were with positive score hence considered for further analysis. All the peptides predicted to be nontoxic by toxipred online toxicity prediction tool.

**Table 1:**
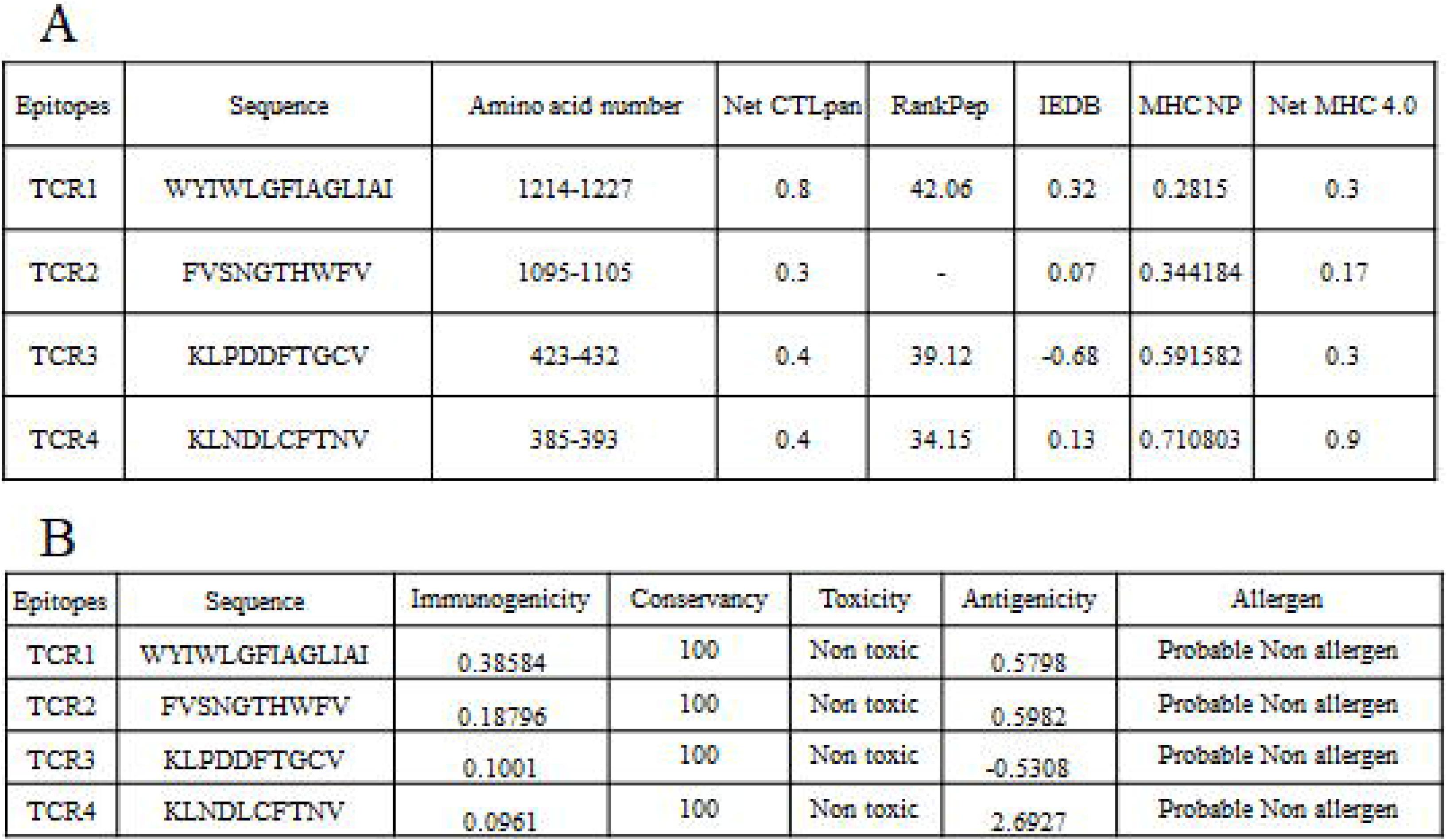
A. Common peptides predicted by at least four TCR epitopes prediction tools; B. immunogenic properties of that peptides. In case of immunogenic properties peptide number 3 has negative score for antigenicity hence neglected for further analysis.

| Epitopes | Sequence | Amino acid number | Net CTLpan | RankPep | IEDB | MHC NP | Net MHC 4.0 |
| --- | --- | --- | --- | --- | --- | --- | --- |
| TCR1 | WYTWLGFIAGLIAI | 1214-1227 | 0.8 | 42.06 | 0.32 | 0.2815 | 0.3 |
| TCR2 | FVSNGTHWFFV | 1095-1105 | 0.3 | - | 0.07 | 0.344184 | 0.17 |
| TCR3 | KLPDDFTGCV | 423-432 | 0.4 | 39.12 | -0.68 | 0.591582 | 0.3 |
| TCR4 | KLNDLCFTNV | 385-393 | 0.4 | 34.15 | 0.13 | 0.710803 | 0.9 |

B
| Epitopes | Sequence | Immunogenicity | Conservancy | Toxicity | Antigenicity | Allergen |
| --- | --- | --- | --- | --- | --- | --- |
| TCR1 | WYTWLGFIAGLIAI | 0.38584 | 100 | Non toxic | 0.5798 | Probable Non allergen |
| TCR2 | FVSNGTHWFFV | 0.18796 | 100 | Non toxic | 0.5982 | Probable Non allergen |
| TCR3 | KLPDDFTGCV | 0.1001 | 100 | Non toxic | -0.5308 | Probable Non allergen |
| TCR4 | KLNDLCFTNV | 0.0961 | 100 | Non toxic | 2.6927 | Probable Non allergen |

#### B cell receptor linear epitopes

After provision of spike protein sequence to the BepiPred server showed average score of 0.407, with 0.695 maximum and 0.163 minimum scores (Fig1). Other properties like physiology of amino acid residues, accessible surface, hydrophilicity, Fexibility and β turns were also determined (Table 2 and Fig1). Peptides obtained in six different methods were arranged according to higher to lower score and 2 % of high score peptides were selected for overlap screening. Regions commonly overlap in at least four prediction methods were selected as potential B epitopes. Regions 250 to 261 and 1246 to 1267 were overlapped regions as per the condition applied. Region 250 to 260 includes BepiPred region 250-260, 251to257 predicted by Karplus-Schulz flexibility and Chau-Fasman β turn prediction methods, and 250 to 257 regions predicted by parker hydrophilicity prediction method. Similarly, region 1246 to 1267 covers region predicted by BepiPred 2.0 (1252-1267), Emini Surface accessible (1257-1262), Kolaskar and Tongoankar method (1247-1254), Chau-Fasman β turn prediction method (1246-1252) and Parker hydrophilicity method (1256-1262).

**Figure1:**
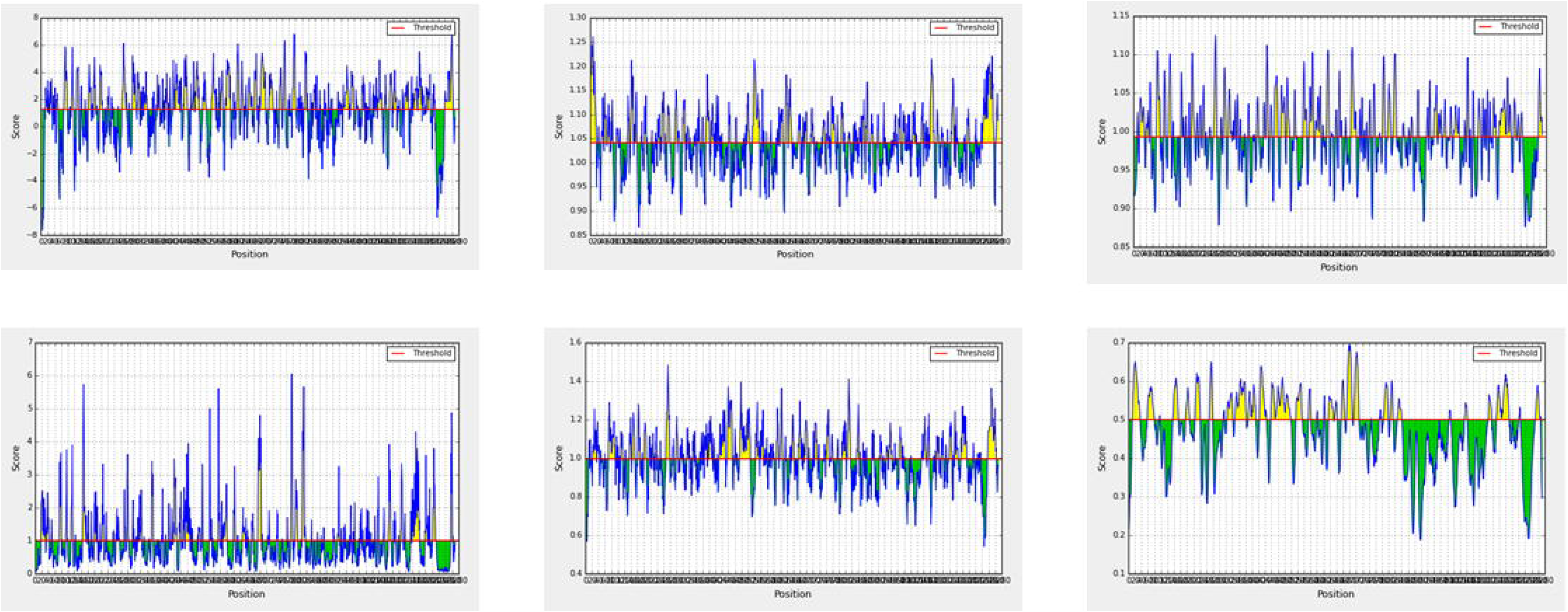
Graphical representation of prediction of B cell epitopes Immunogenic peptide predicted by various tools probable epitopes are scored above the threshold, showed here as yellow. A: Parker hydrophilicity prediction, B: Chau-Fasman β turn prediction C: Karplus-Schulz flexibility, D: Emini Surface accessible, E: Kolaskar and Tongoankar, F: Bepipred Linear Epitope Prediction 2.0

**Table 2:** B cell receptor epitope prediction, A: Common B cell receptor epitopes predicted by various methods B: Score obtained B cell receptor epitopes for Spike glycoprotein of SARS-CoV2

**A**
| Method | Average Score | Maximum Score | Minimum Score |
| --- | --- | --- | --- |
| BepiPred 2.0 | 0.470 | 0.183 | 0.695 |
| Emini Surface accessible | 1.0 | 0.042 | 6.051 |
| Chua-Fasman $\beta$ turn | 0.997 | 0.541 | 1.484 |
| Karplus-Schulz flexibility | 0.993 | 0.876 | 1.125 |
| Kolaskar-Tongaonkar antigenicity | 1.041 | 0.866 | 1.261 |
| Parker hydrophilicity | 1.238 | -7.629 | 7.743 |

**B**
| BCR epitopes | Region | Sequence |
| --- | --- | --- |
| BCR1 | 250-260 (11 aa) | TPGDSSSGWTA |
| BCR2 | 1246-1267 (22 aa) | GCCSCGSCCKFDEDDSEPVLKG |

### Engineering of multi epitope vaccine sequence

Three peptides high scored, predicted for T cell receptors and two peptides for B cell receptors were ligated together by GPGPG or AAY. Additionally, OmpA (GenBank: AFS89615.1) was linked at N terminal end of the designed polypeptides by EAAAK linker. For purification purpose sis histidine residues were added. Final amino acid residues of designed vaccine were 454 when all the five peptides, linkers and adjuvant were ligated (Fig 2).

**Figure 2.**
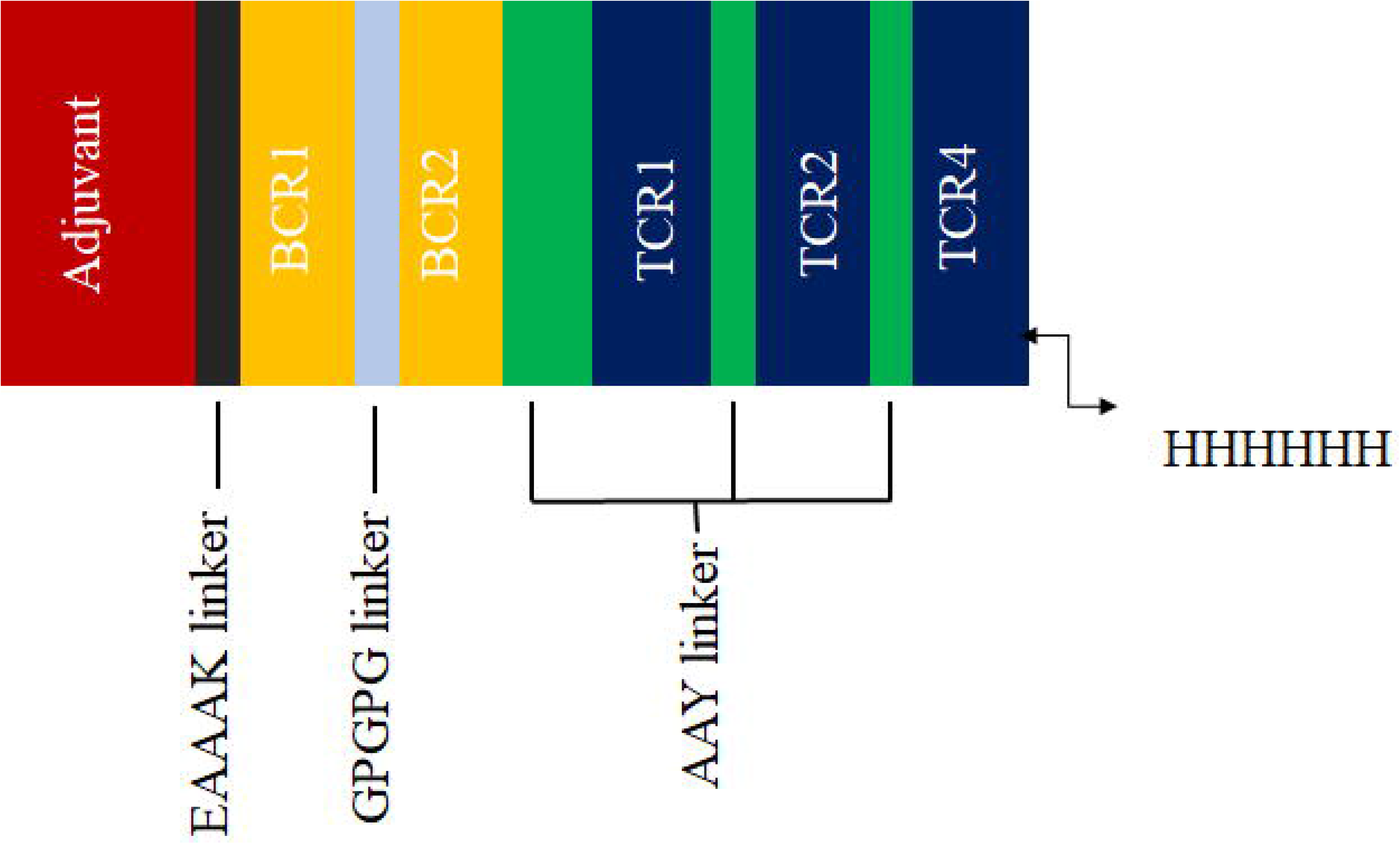
Schematic presentation of the final multi-epitope vaccine peptide. The 599-amino acid long peptide sequence containing an adjuvant (Red) at the amino terminal end linked with the multi-epitope sequence through an EAAAK linker (black). BCR epitopes (yellow) are linked using GPGPG linkers (cyan) while the TCR epitopes (blue) are linked with the help of AAY linkers (green). A 6x-His tag is added at the Carboxy terminus for purification and identification purposes.

### Prediction of immunogenic properties of designed vaccine

Antigenicity of the designed multi-epitopes vaccine predicted by VaxiJen v2.0 server was 0.7048 with probable antigen annotation for virus as target organism at 0.5 threshold. Antigenicity predicted for same sequence of vaccine by AntigenPro server was 0.765946 with 0.953508 predicted probable solubility upon overexpression. Antigenicity of peptide without adjuvant sequence was 0.671028 with 0.861451 probable solubility of peptides upon overexpression. The results obtained indicates that with and without adjuvant predicted vaccine sequence is potentially antigenic in nature. AllergenFP and AllerTop servers predicted that both vaccine with adjuvant and without adjuvant are probable non allergen with 0.84 and 0.77 Tanimoto similarity indexes, respectively.

### Prediction physiochemical properties

The computed molecular weight of designed vaccine was 48.49368 KDa and theoretical pI is 7.96. Predicted pI indicates the slight alkalinity of the vaccine (Table 3). The estimated half-lives were 30 hrs in mammalian reticulocytes (in vitro), >20 hours in yeast (in vivo) and >10 hours in Escherichia coli (*in vivo*). Vaccine is highly stable with instability index 24.58 that classify the protein as stable. Estimated aliphatic index 79.02 indicating thermo-stability of the protein(70). The predicted GRAVY score is −0.175 (Table 3). Negative GRAVY score indicates that protein is hydrophilic in nature and will interact with water molecules(71).

**Table 3.** Analysis of the physicochemical and immunological properties of the designed vaccine for SARS-CoV2

| Characters | score |
| --- | --- |
| Antigenicity (AntigenPro and VaxiJen v2.0) | 0.765946 and 0.7048 (Probable antigen) |
| Allergen (AllergenFP and AllerTop ) | 0.84 and 0.77 (probable non allergen) |
| Possible Solubility | 0.9557226 |
| Number of amino acids | 454 |
| Molecular weight | 48.49368 |
| pI | 7.94 |
| The instability index (II) | 24.58 |
| Aliphatic index | 79.02 |
| Grand average of hydropathicity (GRAVY) | -0.172 |

Secondary structure prediction: Secondary structure predicted by online tool RaptorX property includes 16% alpha helix region, 42% beta sheet region and 40% coil region. Furthermore, solvent accessibility of amino acid residues predicted to be 42% exposed, 27% medium exposed and 24% buried. Total 56 residues (12%) are predicted as residues in disordered region. Pictorial representation of secondary structure predicted of final protein by PSIPRED is shown in figure 3.

**Figure 3:**
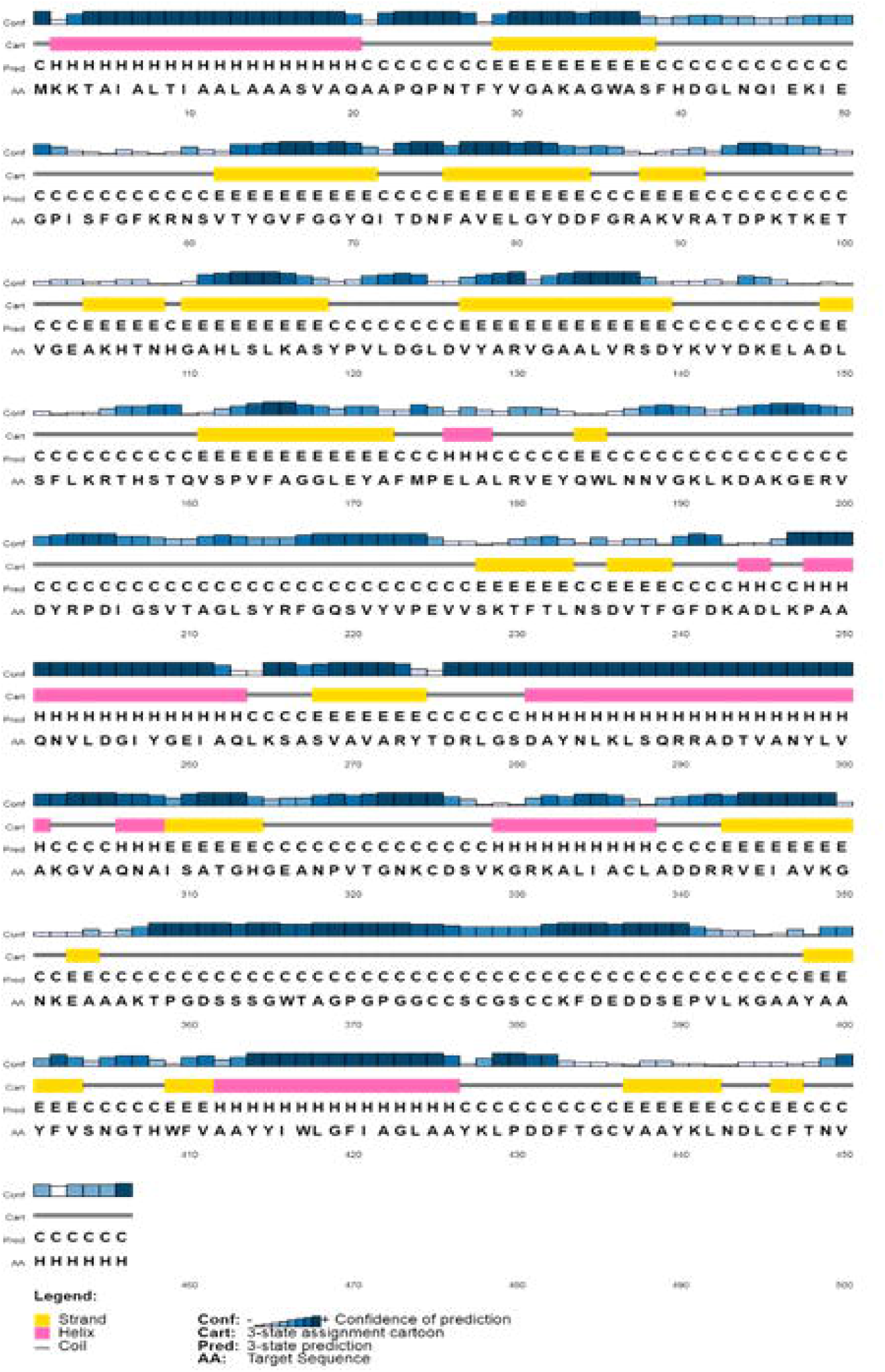
Graphical representation of secondary structure characters; 42% beta sheet region, 40% coil region and 16% alpha helix region

### Tertiary structure prediction refinement and validation

I-TASSER predicted the top five 3D structure model of designed vaccine utilizing 10 threading templates. Z score for these 10 templates was ranging from 0.95 to 5.50 indicating good alignment with sequence submitted. C score is critical in the quality of built model which is quantitatively measured. C score ranges in between −5 to 2 signifies the best model quality. C score of top five model ranging from −4.14 to −3.41. Model with high C score i.e. −3.41, estimated TM score 0.34±0.11 and estimated RMSD 15.6±3.3 was chosen for further analysis (Fig 4A)(72).

**Figure 4:**
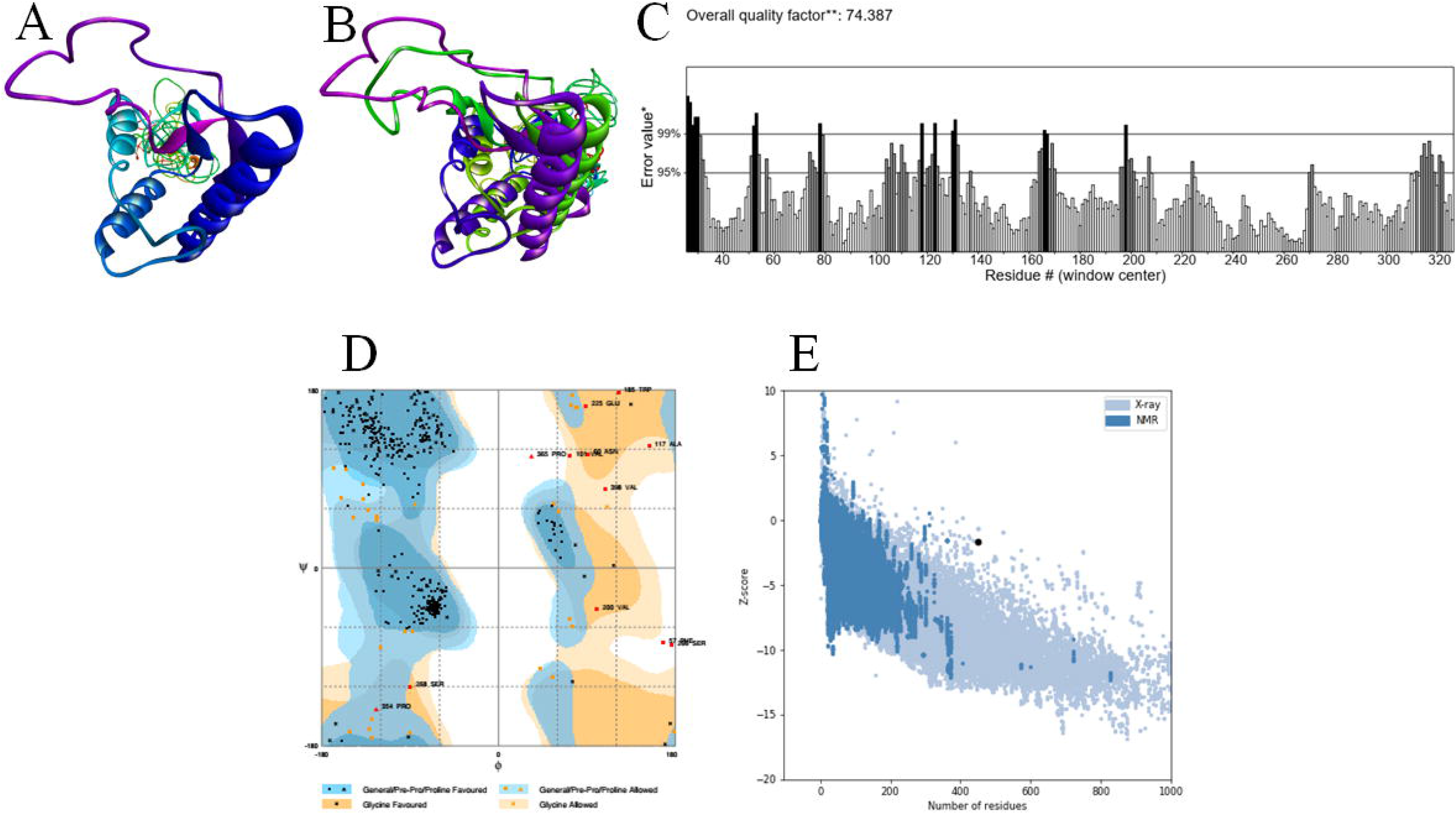
Vaccine tertiary structure modelling, refinement and validation A: tertiary structure of ITASSER modelled multi-epitopes Vaccine along with OmpA as adjuvant; B: Refined 3D model (green) after GalaxyRefine superimposed on ITASSER crude model (violet) C: Errata score for refined model with graphical representation, D: Ramachandran plot for refined 3D model of vaccine, E: Z-score −1.64 by ProSA webtool.

For refinement purpose tertiary structure predicted by I-TASSER is initially submitted to ModRefiner and finally to GalaxyRefine. Among top five refined models, model 5 found to be the best based on various parameters like GDT-HA 0.8398, RMSD-.696, MolProbit-3.526 and Rama favored 77.1(Fig 4B).

Quality of refined model 5 was validated by Ramachandran plot at RAMPAGE webserver. 79.5% residues of modelled vaccine are in favored region, 13.1% in allowed region and 7.3% are in outlier region (Fig 4D). Error in the model are analyzed by ProSA web and ERRAT web server. Z score is −1.64 (Fig 4E) and ERRAT score index is 74.387 (Fig 4C). Ramachandran Plot Z score and ERRAT score showed that refined model 5 is of good quality and can be used further.

### Dis-continuous epitopes for B cell

Four discontinuous epitopes include of 265 residues were predicted. The score of the predicted epitopes ranges from 0.58 to 0.749. Shortest and longest discontinuous B cell epitope is of 14 and 97 residues long respectively.

### Molecular Docking

HADDOCK web bases server has been utilized to dock the designed vaccine to TLR2 and TLR 4. TLR2/4 eliciting immune response toward designed vaccine was analyzed by conformational change with reference to adjuvant TLR2/4 complexes. Top models from each group with lowest HADDOCK score were selected. HADDOCK score for TLR2/Adjuvant and TLR2/Vaccine were, −89.3 and −95.9 Kcal/mol, respectively. In addition, relative binding energies were −9.8 and −12.8 for the TLR2/adjuvant and TLR2/vaccine, respectively. Similarly, HADDOCK scores for TLR4 adjuvant complex and TLR4/vaccine complex were −75.6 and −139.9 Kcal/mol, respectively. Relative binding energy of TLR4/adjuvant complex is −10.7 Kcal/mol and −8.2 for TLR4 vaccine complex. Difference in scores and binding energies of adjuvants and vaccines indicates the change in conformation that may stimulate TLR2/4 receptors. Even there is difference in number interacting residues at the juncture. TLR2/Adjuvant complex have charged-charged 2, charged polar 6, charged-apolar 24, polar-polar 2, polar-apolar 22 and apolar-apolar21. While TLR2/Vaccine have charged-charged 3, charged polar 7, charged-apolar 15, polar-polar 1, polar-apolar 13 and apolar-apolar 119(Fig 5A, B). Similarly, TLR4/Adjuvant complex have charged-charged 3, charged polar 10, charged-apolar 21, polar-polar 3, polar-apolar 15 and apolar-apolar20. While TLR4/vaccine complex has charged-charged 1, charged-polar 9, charged-apolar 10, polar-polar 9, polar-apolar 15 and apolar-apolar 8 interface contacts were observed (Figure 5C, D).

**Figure 5:**
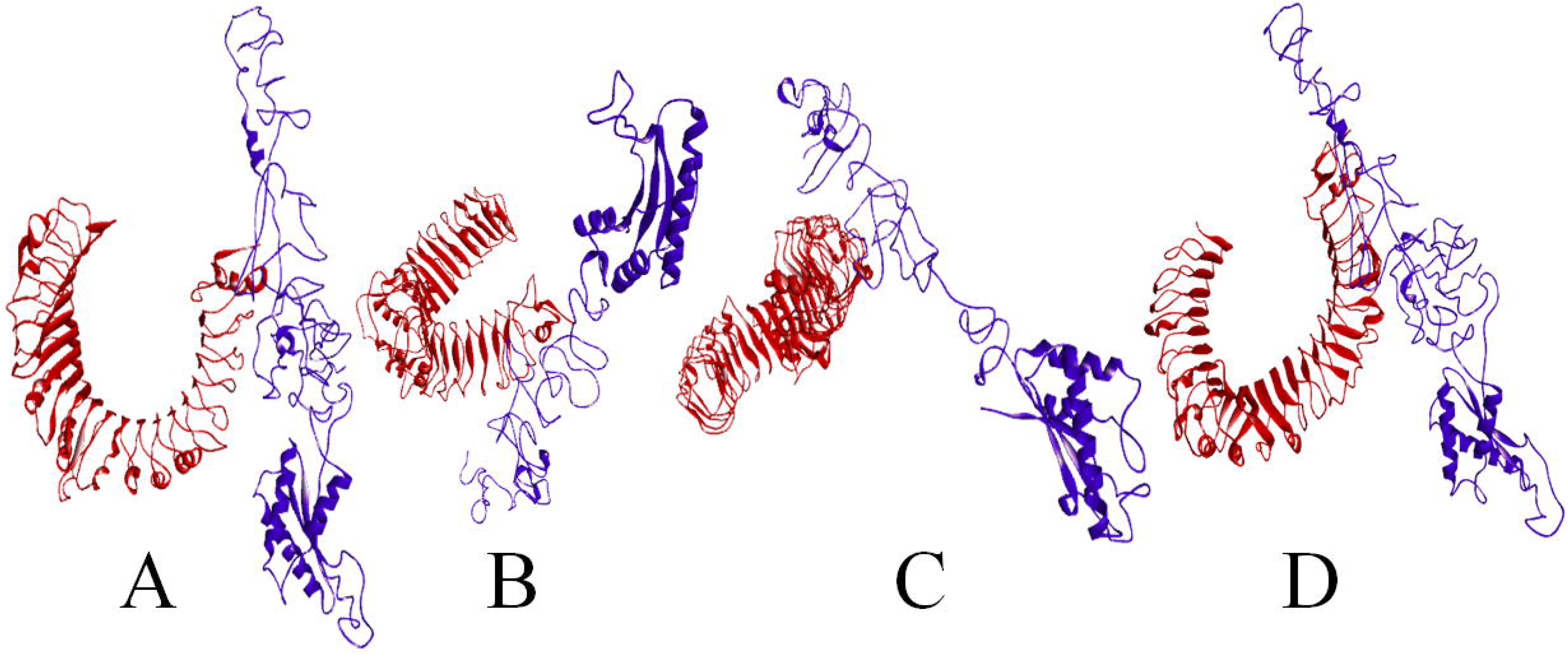
Haddock docking cartoon models showing interaction A: TLR2-Vaccine; TLR2 in red and Vaccine in blue; B:TLR2-Adjuvant; TLR2 in red and Adjuvant in blue; C: TLR4-Vaccine; TLR4 in red and Vaccine in red; D: TLR4-Adjuvant; TLR4 in red and Adjuvant in blue

### Codon optimization

Enhance expression of nucleotide construct in Escherichia coli, is reverse translated in Jcat online webtool. Total number of nucleotides in the constructs were1350. After codon optimization CAI was 1.0 and average GC content of optimized nucleotide construct was 50.7340 indicating good probable expression of vaccine in *E. coli* K12. To clone the gene Xho and XbaI restriction sites are added at N and C terminus of the construct and gene was cloned in pET30a+ plasmid, by using SnapGene software (Fig 6).

**Figure 6:**
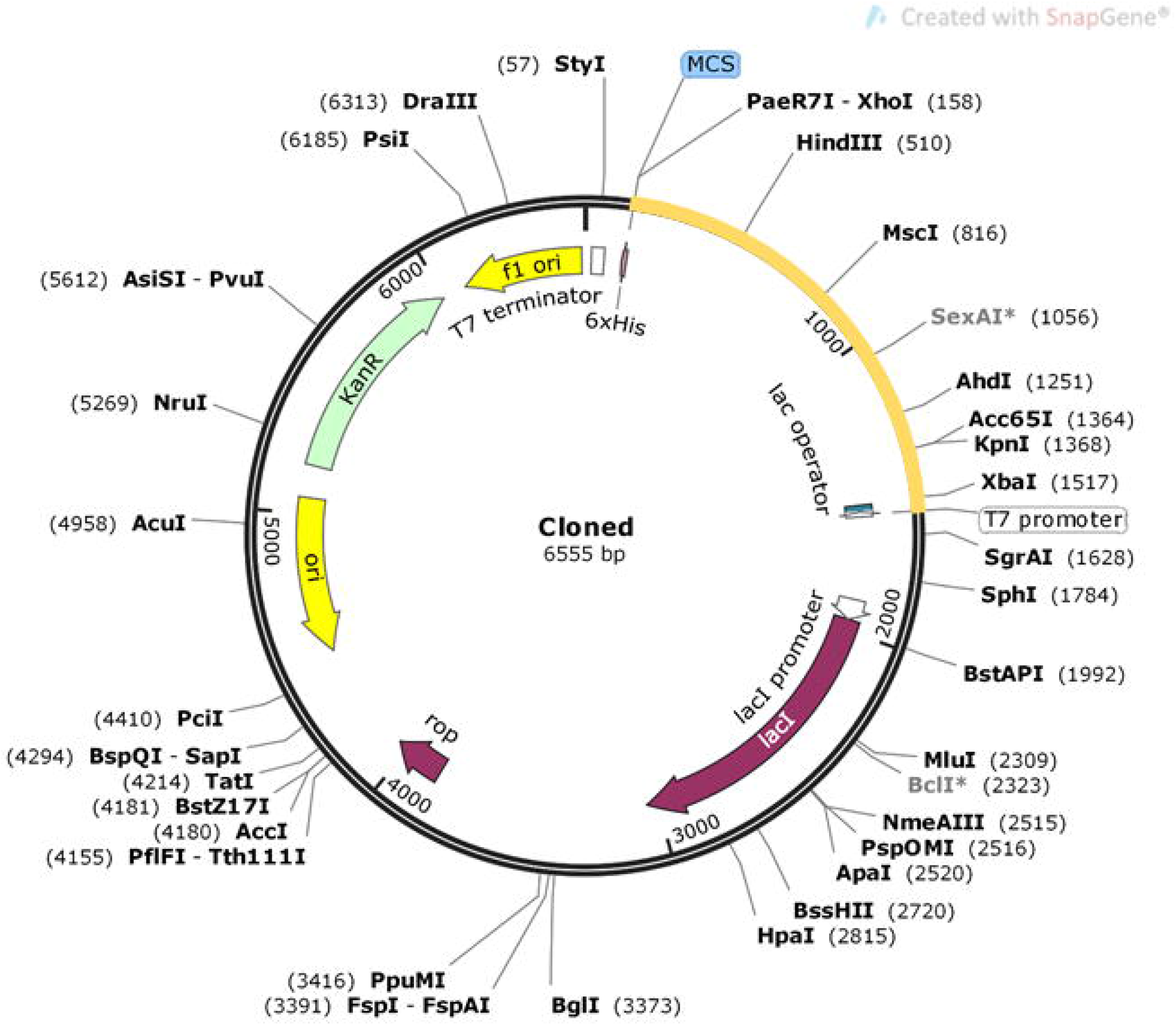
*In silico* cloning of designed vaccine construct in pET30a+ expression vector where yellow arc segment represents cloned construct and remaining segment represent vector backbone.

**Figure 7:**
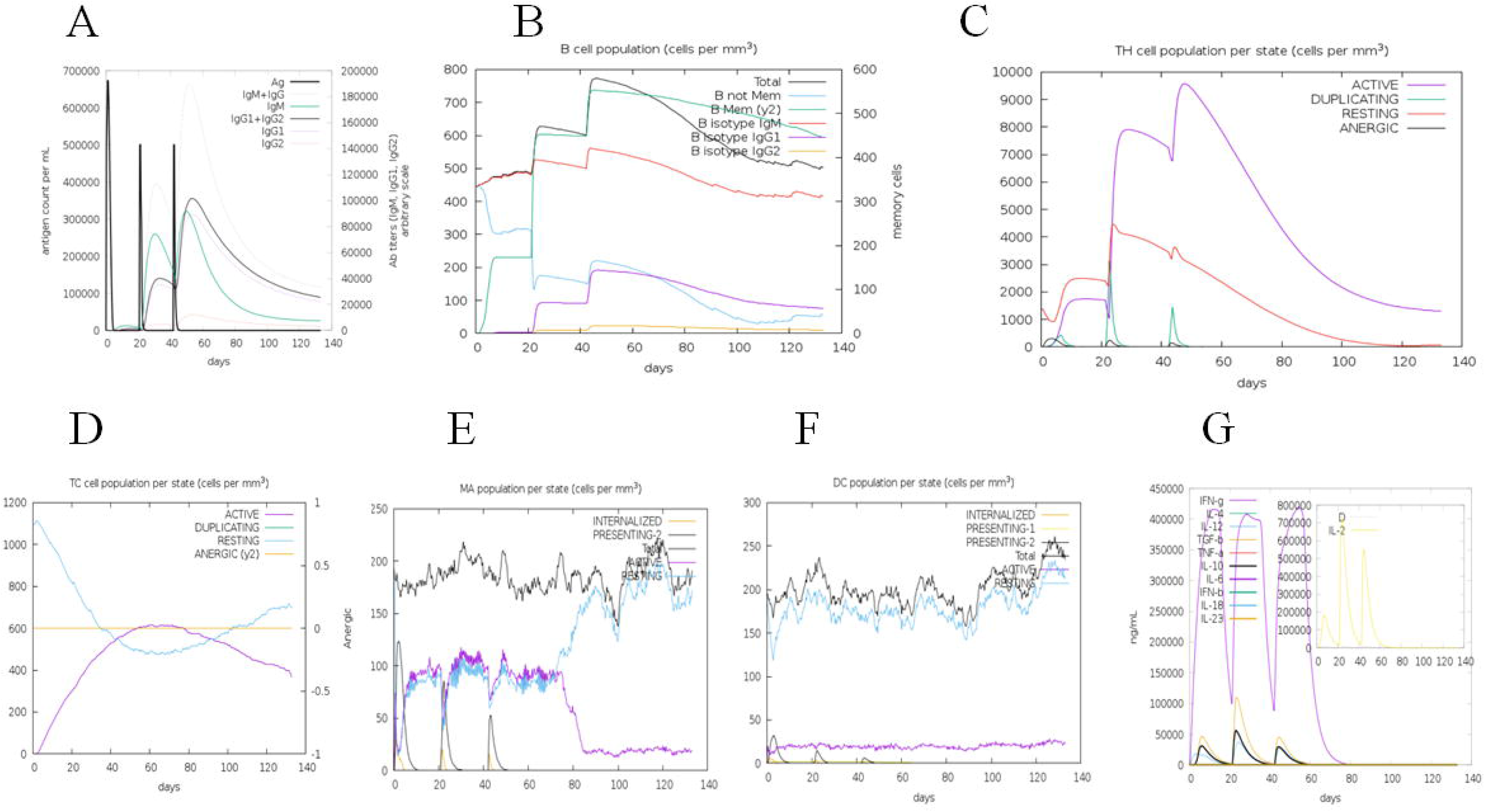
Immune response simulation by designed vaccine, A: Immune response upon antigen exposure; B: B cell population per state(cells/mm3); C: TH cell population state (cells/mm3); D: TC cell population state (cells/mm3); E: activity of macrophage population in three subsequent immune responses; F: Dendritic cell population state (cells/mm3) upon antigen exposure; G:Cytokine induced by three injection of vaccine after one week interval, The main plot shows cytokine levels after the injections. The insert plot shows IL-2 level with the Simpson index, D indicated by the dotted line

### Immune simulation

C-ImmSim Immune simulator web server was used for determining ability of vaccine to induce immunity. Results obtained are consistent with the experimental results published elsewhere(73). Within first week of vaccine administration primary response was observed by high level of IgM. Secondary and tertiary immune response clearly seen with increase in B cell number, level of IgM, IgG1+IgG2, and IgM+IgG with decreasing concentration of antigen. Different B cell isotypes population were also found increasing indicating isotype switching and memory formation (Fig 6A). B cell activities were also found high, especially B Isotype IgM and IgG1, was observed with prominent memory cell formation (Fig 6B). Similarly, cell population of Th and Tc cells are found high along with memory development (Fig 6C, D). In addition, macrophage active was found to be increased consistently after each antigen shots and declined upon antigen clearance (Fig 6E). Another cell from cell mediated immunity, dendritic cells were also found increased (Fig 6F). IFNγ and IL2 expression was found high with low Simpson Index indicating sufficient immunoglobulin production, suggesting good humoral immune response (Fig 6G). Simpson Index (D) is a measure of diversity. Increase in D over time indicates emergence of different epitope-specific dominant clones of T-cells. The smaller the D value, the lower the diversity.

## Discussion

Till date there is no cure available for COVID19, suggesting urgent requirement of drug or vaccine to control the spread of SAR-CoV2 infections. Advancement in bioinformatics tools, process of vaccine development can be facilitated by identifying potential epitopes for T and B cells that will lead to vaccine for prevention of COVID19. Several reports and animal trials suggest that S-protein can be potential target for vaccine development(22,73–76). There are several reports suggesting the importance of Spike protein in activation of cells of immunity and vaccine development(77–79). HLA-A2 restricted epitopes from S protein of SARS-CoV2, elicit T cell specific immune response in SARS-CoV2 recovered patients(27). Similarly serum samples of SARS-CoV2 recovered patients were sufficient to neutralize epitope rich region on the spike S2 protein from SARS-CoV2 (80). Programing of live attenuated form of pathogen is commonly used strategy for the vaccine development. Despite efficacy of such vaccines, reversion of virulence remains a challenge and not utilized in weak immune patients(81,82). Here in case SARS-CoV2 due to its high communal transmission vaccine research requires highly skilled workers, sophisticated instrumentation and biosafety level III facility. This may likely the slow down the process of vaccine development, enhanced cost leading to restriction on availability of vaccine for mass population. Advancement in the field of bioinformatics and molecular biology techniques have provided opportunity of development of vaccine with high efficacy, less time for development, low cost of production that may lead to low cost vaccine in time for large population. We designed a multi-epitopes vaccine construct from S-protein of SARS-CoV2. Several online tools were used for the designing of epitopes and thereby a vaccine. From various epitopes predicted by the online server based on common sequence and high score three TCR and two BCR epitopes were selected as part of COVID19 vaccine. All the five TCR and BCR epitopes were linked by GPGPG and AAY linker peptides (Fig 2). To enhance the antigenicity of TCR/BCR linked epitopes peptide, adjuvant OmpA were linked by EAAAK linker sequence for a complete vaccine design which was found to restore antigenicity and non-allergen status. (Table 2). For successful vaccine candidate, secondary and tertiary structures characters play an essential role. Secondary structure of designed vaccine contains 16% alpha helixes, 42% beta sheets and 40% coils. Upon refinement tertiary structure of designed vaccine showed improvement to desirable level as indicated by Ramachandran plot. Designed vaccine showed high antigenicity score, no probable allergen.

TLR are essential receptor proteins in activation of innate immune response. That recognize and respond by induction of immune reactions to PAMPs (pathogen associated molecular patterns). Various TLRs have shown to activate immune response to virus through their interaction with nucleic acids and envelopes proteins. TLR3 and TLR7/8 involved in nucleic acids detection, while TLR2 and TLR4 recognize envelope glycoproteins(83). Interaction of S protein with TLR2 increase the production of IL8 in human monocyte macrophages(84). TLR 2 also involved in eliciting innate immune response upon recognition of several other virus components(85,86). In a comparative binding efficacy of S-proteins to TLRs, TLR 4 showed strongest protein-protein interaction with hydrogen and hydrophobic bonds on extracellular domain(87). In molecular docking analysis, TLR2 and TLR4 showed stable protein-protein interaction with designed vaccine compared to adjuvant protein. TLR 2 showed more stable interaction than TLR4, which is consistent with previous report(87). Stable interaction of designed vaccine indicates efficiency of vaccine for activation of TLRs, involved in dendritic cell activation, thereby subsequent antigen processing, and presentation on surface to T cells(88). Immune simulation showed typical natural immune response pattern upon multiple exposure to same antigen. Server predicted elevated level of B cell and T cell for longer time on repeated exposure of antigen. Increased level of antiviral cytokine IFNγ and IL2 indicate potential of subsequent activation of T-helper cell thereby high level of Ig production, supporting humoral immune response(89).

To validate the designed vaccine in the screening of immune reactivity by serological test, it is necessary to express the construct in preferred host like *E. coli* K12 for recombinant proteins. Codon optimization of was performed to achieve high expression designed vaccine in E coli 12. Codon adaptability index (CAI=1.0) and GC content (51.66%) of the vaccine construct was optimum for high expression of recombinant protein in host. Immediate step, here after is to express the protein in preferred host and validate results obtained in this report by performing the several immunological assays.

## Conclusion

Inadequate number of drugs and time-consuming process of drug development, multiplication chain of SARS-CoV2 infection can be control only by vaccine. We utilized tools and techniques of immune informatics to design a potential vaccine candidate. This vaccine codes epitopes form S protein of SARS-CoV2 virus for T and B cell receptors. This is one more time showed that bioinformatics approach can effectively use to develop vaccines in short time and with incurred less cost. Although *in silico* results point out the effectiveness of the vaccine, efficacy need to be analyzed by preforming laboratory experiments and animal model studies.

